# Caskin2 is a novel talin and Abi1-binding protein that promotes cell motility

**DOI:** 10.1101/2024.03.15.585234

**Authors:** Wei Wang, Paul Atherton, Maaike Kreft, Lisa te Molder, Sabine van der Poel, Liesbeth Hoekman, Patrick Celie, Robbie P. Joosten, Reinhard Fässler, Anastassis Perrakis, Arnoud Sonnenberg

**Affiliations:** Division of Cell Biology, The Netherlands Cancer Institute, Plesmanlaan 121, Amsterdam 1066 CX, The Netherlands; Present address: Shanghai Cancer Institute, Renji Hospital, Shanghai Jiao Tong University School of Medicine, Dongfang Road, Shanghai 200127, China; Department of Molecular and Clinical Cancer Medicine, Institute of Systems, Molecular and Integrative Biology, The University of Liverpool, Liverpool, L69 7BE, UK; Proteomics Facility, The Netherlands Cancer Institute; Oncode Institute and Division of Biochemistry, The Netherlands Cancer Institute; Department of Molecular Medicine, Max Planck Institute of Biochemistry; Division of Biochemistry, The Netherlands Cancer Institute

**Author notes:** Correspondence and request for materials should be addressed to A.S. W. Wang and P. Atherton contributed equally.

**Keywords:** Caskin2, talin, focal adhesion, KANK, cortical microtubule stabilizing complex, Abi, WAVE

## Abstract

Talin couples the actomyosin cytoskeleton to integrins and transmits tension to the extracellular matrix. Talin also interacts with numerous additional proteins capable of modulating the actin-integrin linkage and thus downstream mechanosignaling cascades. Here, we demonstrate that the scaffold protein Caskin2 interacts directly with the R8 domain of talin through its C-terminal LD motif. Caskin2 also associates with the WAVE Regulatory Complex to promote cell migration in an Abi1-dependent manner. Furthermore, we demonstrate that the Caskin2-Abi1 interaction is regulated by growth factor-induced phosphorylation of Caskin2 on serine 878. In MCF7 and UACC893 cells, which contain an amplification of *CASKIN2*, Caskin2 localizes in plasma membrane-associated plaques and around focal adhesions in CMSCs. Taken together, our results identify Caskin2 as a novel talin-binding protein that may not only connect integrin-mediated adhesion to actin polymerization, but could also play a role in crosstalk between integrins and microtubules.

## Introduction

Cell migration is essential for several biological processes including embryogenesis, wound repair, and tissue homeostasis, and also underpins metastasis of cancer cells. Migrating cells form actin-driven protrusions that are linked to the underlying extracellular matrix (ECM) by integrin-mediated adhesions. Activated integrins, which engage with ECM components, dynamically cluster at the leading edge. The assembly/disassembly of integrin-mediated adhesions is regulated by actin polymerization; furthermore, many proteins capable of influencing actin polymerization are known to bind to integrin-associated proteins. Thus, integrins and polymerizing actin are linked by complex feedback systems that remain poorly defined.

Actin polymerization at the leading edge is driven by the Arp2/3 complex, which is regulated by a complex of five proteins (WASF1/2, CYFIP1/2, ABI1/2, NCKAP1, HSPC300) called the WAVE Regulatory Complex (WRC) (Rottner et al., 2021). Meanwhile, integrins are indirectly connected to actin via multidomain proteins including talin and vinculin, that undergo force-mediated conformational changes that affect their dynamics and the recruitment of additional proteins (Jansen et al., 2017). Talin cycles between the cytosol, where it remains auto-inhibited, and the plasma membrane, where it binds integrin tails upon activation (Goult et al., 2013a). The N-terminal FERM domain of talin binds integrins, PIP_2_, Arp2/3, and F-actin (actin binding site 1; ABS1), whilst the long C-terminal rod domain consisting of 13 alpha-helical bundles (R1-R13) (Calderwood et al., 2013) contains two additional actin binding sites (ABS2 and ABS3 (Atherton et al., 2015)) and 11 vinculin binding sites (Calderwood et al., 2013) that are exposed upon force-mediated unfolding of the talin rod domains (Goult et al., 2018; Yao et al., 2016). In addition to integrin, actin and vinculin, multiple other proteins have been identified to interact with talin, including paxillin and DLC1 (Zacharchenko et al., 2016), α-synemin (Sun et al., 2008), tensin3 (Atherton et al., 2022), KANK-family members (Bouchet et al., 2016; Sun et al., 2016), RIAM (Lee et al., 2009), and CDK1 (Gough et al., 2021). The interaction of KANK family proteins with talin has recently attracted considerable attention (Bouchet et al., 2016; Guo et al., 2023; Li et al., 2023; Rafiq et al., 2019; Sun et al., 2016) as the KANK proteins can connect to microtubules through a complex of proteins, referred to as the cortical microtubules stabilization complex (CMSC; (Noordstra and Akhmanova, 2017)), which localizes at the plasma membrane. These CMSCs are strongly clustered around focal adhesions at the leading edge and may play an important role in their turnover (Rafiq et al., 2019; Sun et al., 2016).

Here, we identified Caskin2 as a novel component of CMSCs and an interactor of both talin and Abi1. The talin interaction is mediated by a C-terminal LD-motif, whilst Abi1 binding occurs via the Abi1-SH3 domain interacting with the proline-rich region of Caskin2. Furthermore, we show that the Caskin2-Abi1 interaction is regulated by phosphorylation of S878 of Caskin2, which occurs downstream of EGFR and MEK1/2 activation. Meanwhile, over-expression of Caskin2 promotes cell motility. Finally, we show that in Caskin2 is a component of the CMSCs. Overall, our results identify Caskin2 as a scaffold protein connecting integrin-mediated adhesions to actin polymerization and microtubules.

## Results

### Caskin2 is a scaffold protein in proximity to cell-ECM adhesion sites

We previously identified Caskin2 (*CASKIN2*) in proximity of integrin α6β4 in PA-JEB/β4 keratinocytes (β4-deficient keratinocytes reconstituted with wild-type β4) by a proximity-dependent biotin identification (BioID) assay combined with mass spectrometry-based proteomics (Te Molder et al., 2020). Caskin2 is a relatively unexplored scaffold protein composed of six ankyrin repeats (ANK), an atypical (Kwan and Donaldson, 2016) Src homology 3 (SH3) domain and tandem sterile α-motifs (SAM) domains, followed by an extended disordered proline-rich region (PRR) and a conserved C-terminal domain (CTD) (Marchler-Bauer et al., 2015; Smirnova et al., 2016) (Fig. 1A). Caskin2 is widely expressed in adult tissues, whereas expression of Caskin1 is restricted to the brain (Human Protein Atlas). Whilst the majority of studies on Caskin2 have focused on its role in neuronal cells, its interaction partners, localization and biological function in non-neuronal cells remain unexplored.

**Fig. 1.**
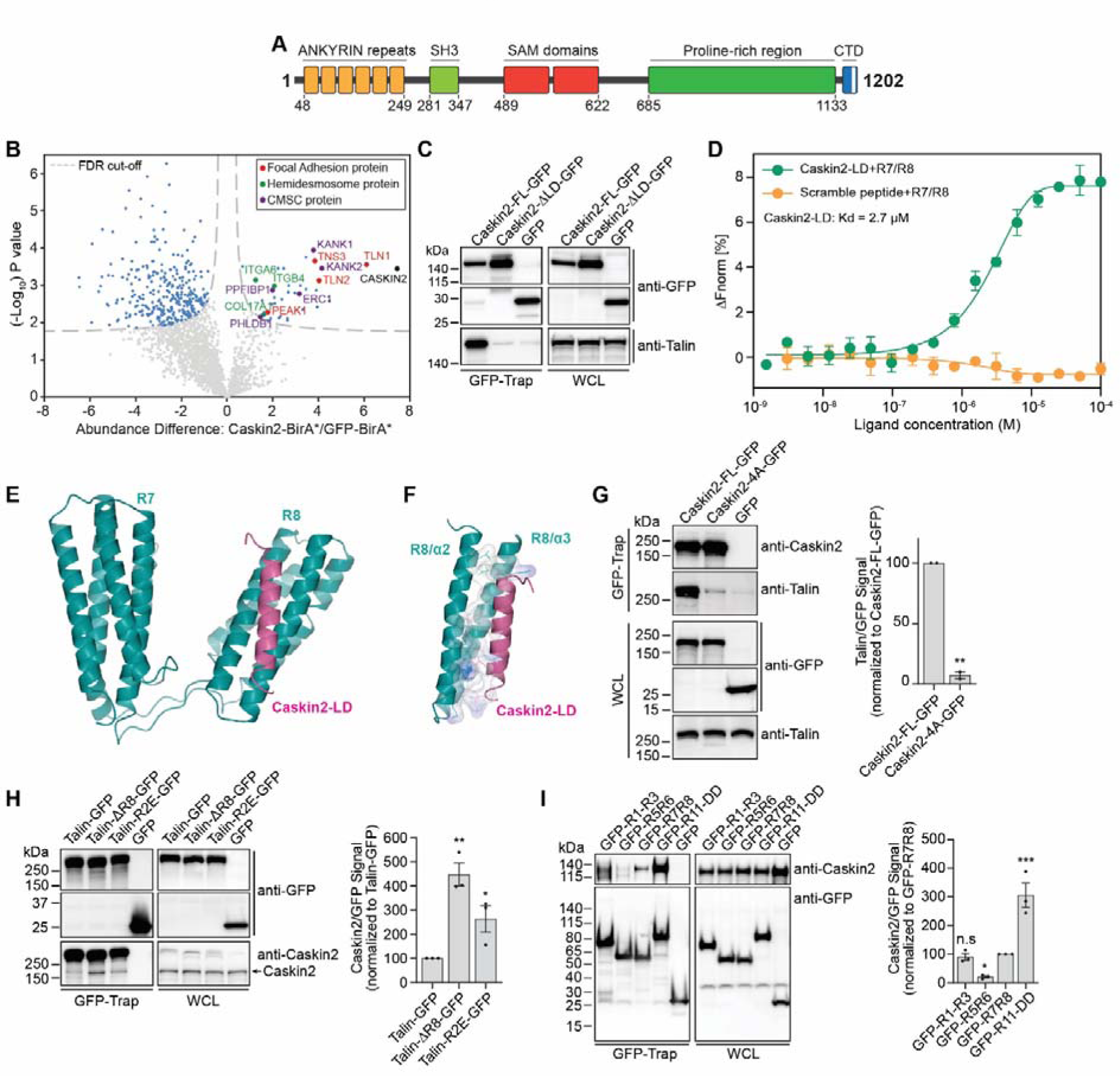
Caskin2 binds directly to talin via a C-terminal LD motif. (***A***) Cartoon of Caskin2; the C-terminal domain contains an LD-motif (white bar). (***B***) Proximity biotinylation assays were performed with PA-JEB/β4 keratinocytes ectopically-expressing either Caskin2 fused to the biotin ligase BirA*, or GFP-BirA* as a negative control. The volcano plot shows the results from three independent experiments (threshold FDR: 0.01 and S0: 0.1). Significant proximity interactors of Caskin2 and GFP are indicated in light blue (GFP interactors, left and Caskin2 interactors, right), red (FA proteins), green (HD proteins) or purple (CMSC components). (***C***) GFP-trap pull-down of Caskin2-GFP and Caskin2-ΔLD-GFP expressed in GE11 ^tetON^ ^β1^ cells, followed by Western blotting for talin and Caskin2. (***D***) MST assay demonstrating binding of Caskin2-LD peptide to the talin R7/R8 peptide (n≥5). Peptide with scrambled amino-acid sequences was used as negative control. Error bars show standard deviation. (***E***) X-ray structure of talin R7R8 (residues 1358-1653, cyan) fragment in complex with Caskin2-LD peptide (residues 1187-1202, magenta). (***F***) Structure analysis show the hydrophobic interaction (with surface charge distribution) of Caskin2-LD (magenta ribbon) with talin R8 (cyan ribbons). Residues of both Talin R8 and Caskin2-LD peptide that form the interface between the two molecules are shown in sticks. The electrostatic surface of these residues are shown (blue for positive, red for negative (not present) and white for neutral). (***G***) GFP-Trap pull-down of GFP, Caskin2-GFP and Caskin2-4A-GFP expressed in COS-7 cells. Proteins in the GFP-trap sample and WCL (input) are detected by Western blotting, using antibodies against talin, Caskin2 and GFP. (***H*** *and **I***) GFP-Trap pull-down in COS-7 cells expressing Caskin2 (without a GFP tag) together with talin-GFP, talin-ΔR8-GFP and talin-R1523E/K1530E-GFP (***H***) or the indicated GFP-tagged talin polypeptides (***I***). Proteins in the GFP-trap sample and WCL (input) are detected by Western blotting, using antibodies against Caskin2 and GFP. Error bars indicate standard deviation; * indicates p<0.05, ** indicates p<0.01, ***indicates p<0.001, One-Way ANOVA with Uncorrected Fisher’s LSD multiple comparisons test. Western blots in C, G, H and I are representative of 3 independent experiments.

To gain a better understanding of the function and localization of Caskin2, we first used BioID in combination with mass spectrometry to identify Caskin2 interacting proteins. For these experiments, we used PA-JEB/β4 keratinocytes since it was in these cells that we originally identified Caskin2 as a proximal protein of α6β4. After stable expression of Caskin2 fused to the promiscuous biotin ligase BirA* (Caskin2-BirA) in PA-JEB/β4 keratinocytes, biotinylated proteins were captured with streptavidin beads and subjected to mass spectrometry. A total of 36 Caskin2 interacting and proximal proteins were identified, including several components of FAs (talin1, 2 (Klapholz and Brown, 2017), tensin3 (Atherton et al., 2022), PEAK1 (Zuidema et al., 2022a)), hemidesmosomes (ITGA6, ITGB4 and COL17A1 (te Molder et al., 2021)) and cortical microtubule stabilizing complexes (KANK1,2, PPFIBP1 (Liprin-β1), PHLDB1 (LL5α) and ERC1/ELKS (Noordstra and Akhmanova, 2017)) (Fig. 1B; Table S1). Together these data suggest that Caskin2 is in close proximity to different matrix adhesion sites and associated complexes.

We confirmed the subcellular localization of Caskin2 by expressing Caskin2 tagged at the C-terminus with GFP, in PA-JEB/β4 keratinocytes. Confocal microscopy revealed that Caskin2-GFP partially co-localized with both integrin β4 and talin (Fig. S1A), a FA protein that is also a proximity interactor of integrin α6β4 in keratinocytes (Te Molder et al., 2020).

### Caskin2 binds directly to talin via a C-terminal LD-motif

Several proteins interact with talin via leucine-aspartic acid motifs (LD-motifs): short helical motifs that facilitate protein-protein interactions with the helical bundles of the talin rod domains, with the talin R7R8 domains being the best characterized LD-binding domains in talin (Zacharchenko et al., 2016) (Fig. S1B). Both Caskin1 and Caskin2 contain an LD-motif in their conserved C-terminus (Fig. S1C), suggesting they may be binding partners of talin-R7R8. To explore whether i) this interaction occurs in cells and ii) whether the interaction is dependent on the presence of integrin β1, we stably expressed either Caskin2-FL-GFP or a Caskin2 construct with the LD-motif deleted fused to GFP (Caskin2-ΔLD-GFP; L1183-D1202 deleted) in GE11 cells expressing integrin β1 under control of doxycycline (referred to hereon in as GE11 ^tetON^ ^β1^ cells). GFP-Trap experiments in GE11 ^tetON^ ^β1^ cells showed that the LD motif is required for the interaction with talin (Fig. 1C, S1D), which occurred in both the presence and absence of doxycycline, indicating the interaction is independent of integrin β1 (Fig. S1E).

Using microscale thermophoresis (MST) we measured the affinity of the Caskin2 LD-peptide to talin R7R8 as ∼2.7 μM (Fig. 1D). To further understand the mode of this interaction, we crystallized the complex and determined the crystallographic structure at 2.7Å resolution (Fig. 1E). The electron density map showed clear binding of the Caskin2-LD peptide to the R8 motif. The peptide was modelled, and the structure was refined to a R_free_ of 26.9% with excellent geometry (Table S2). The Caskin2-LD peptide adopts a helical conformation that packs against two adjacent helices (α2 and α3) on the surface of the talin-R8 four-helix bundle (Fig. 1F), essentially forming a five-helix bundle. Analysis of the interaction interface (Krissinel and Henrick, 2007) showed that the buried interface area is 790Å. The interaction is formed solely by van der Waals contacts between the talin-R8 and caskin2-LD helices, as hydrogen bonds are absent. The Caskin2 residues with the largest buried surface area are L1183 (120Å), F1190 (113Å), L1197 (105Å) and L1201 (104Å). To validate this binding mode, we transiently expressed a GFP-fused Caskin2 mutant with these four amino acids substituted by alanine (termed Caskin2-4A-GFP, Fig. S1F) in COS7 cells and performed GFP-Trap pull down experiments. In contrast to the WT Caskin2-GFP construct, Caskin2-4A-GFP failed to pull-down endogenous talin (Fig. 1G), confirming the interaction observed in the crystal structure.

Our structure of the binding of the Caskin2-LD peptide to talin R7R8 is very similar to the structure of the DLC1-LD peptide binding to talin R7R8 (PDB: 5fzt; (Zacharchenko et al., 2016)), with both forming a five-helix bundle with the R8 domain (Fig. S1G). Interestingly, the relative orientation of the R7 and R8 domains are different in the two structures, demonstrating the structural plasticity between the two talin repeats (Fig. S1H).

### Talin contains multiple Caskin2 binding sites

To investigate whether R8 is the only talin repeat recognized by the LD motif of Caskin2, we expressed GFP-tagged talin lacking the R8 domain (Talin-ΔR8-GFP) together with Caskin2 in COS-7 cells and performed GFP-Trap Co-IP experiments. We also included in this analysis a talin mutant carrying two reverse charge mutations (R1523E and K1530E; Talin-R2E-GFP), which have previously been reported to destabilize R8 (Zacharchenko et al., 2016) and to disrupt binding of the LD-motif containing protein DLC1 (Zacharchenko et al., 2016). Whereas Caskin2 showed only limited binding to full-length exogenous talin (due to the presence of endogenous talin in the COS-7 cell lysates), a slightly increased binding was observed for the charge-reversal talin mutant (R1523E/K1530E), Interestingly, the strongest binding was observed for the talin ΔR8 construct (Fig. 1H). We hypothesize that the increased binding of Caskin2 to the talin ΔR8 construct may arise due to a conformational change in the talin molecule, potentially caused by disruption of the F3-R9 autoinhibition (Goult et al., 2013a) as a result of the deletion of R8, which could expose additional Caskin2 binding sites.

These findings suggest that Caskin2 is able to interact with additional talin rod domains. To provide support for this hypothesis, we used AlphaFold (Evans et al., 2022; Jumper et al., 2021) in the context of the CollabFold implementation (Mirdita et al., 2022) to model the complex between the Caskin2 LD peptide and each individual talin rod domains. This approach suggested a high-confidence complex between the peptide and the R8 domain (closely resembling our crystal structure) and a very low-confidence complex with the R7 domain, which does not bind the peptide in our structure (Fig. S2). The models also suggest that the Caskin2 LD peptide could likely form a complex with the R2, R3, R11, and R12 talin rod domains. To confirm these interactions, we used GFP-tagged talin polypeptides consisting of the R1-R3 domains (GFP-R1-R3), R5 and R6 (GFP-R5R6), R7 and R8 (GFP-R7R8), and R11-R13 including the C-terminal dimerization domain (GFP-R11-DD). GFP-Trap Co-IP experiments using these constructs together with co-expressed Caskin2 in COS-7 cells confirmed the predictions of the AlphaFold analysis, with binding observed between all of the polypeptides and Caskin2 except for GFP-R5R6 (Fig. 1I).

### Caskin2 co-localizes with members of the cortical microtubule stabilizing complex

We next examined the expression levels of endogenous Caskin2 in a variety of human cell types, including HaCaT cells, ovarian carcinoma (OVCAR-4), colorectal carcinoma (HT29, SW480), lung adenocarcinoma (A549), breast cancer (CAMA-1, MCF7, UACC893, MDA-MB-361, MDA-MB-453, T47D) and hepatocellular carcinoma (Hep 3B, Hep 2G). We found high levels of Caskin2 expression in MCF7 and UACC893 breast cancer cells (Fig. 2A), in agreement with previous reports that these cells contain an amplification of *CASKIN2* (Shadeo and Lam, 2006).

**Fig. 2.**
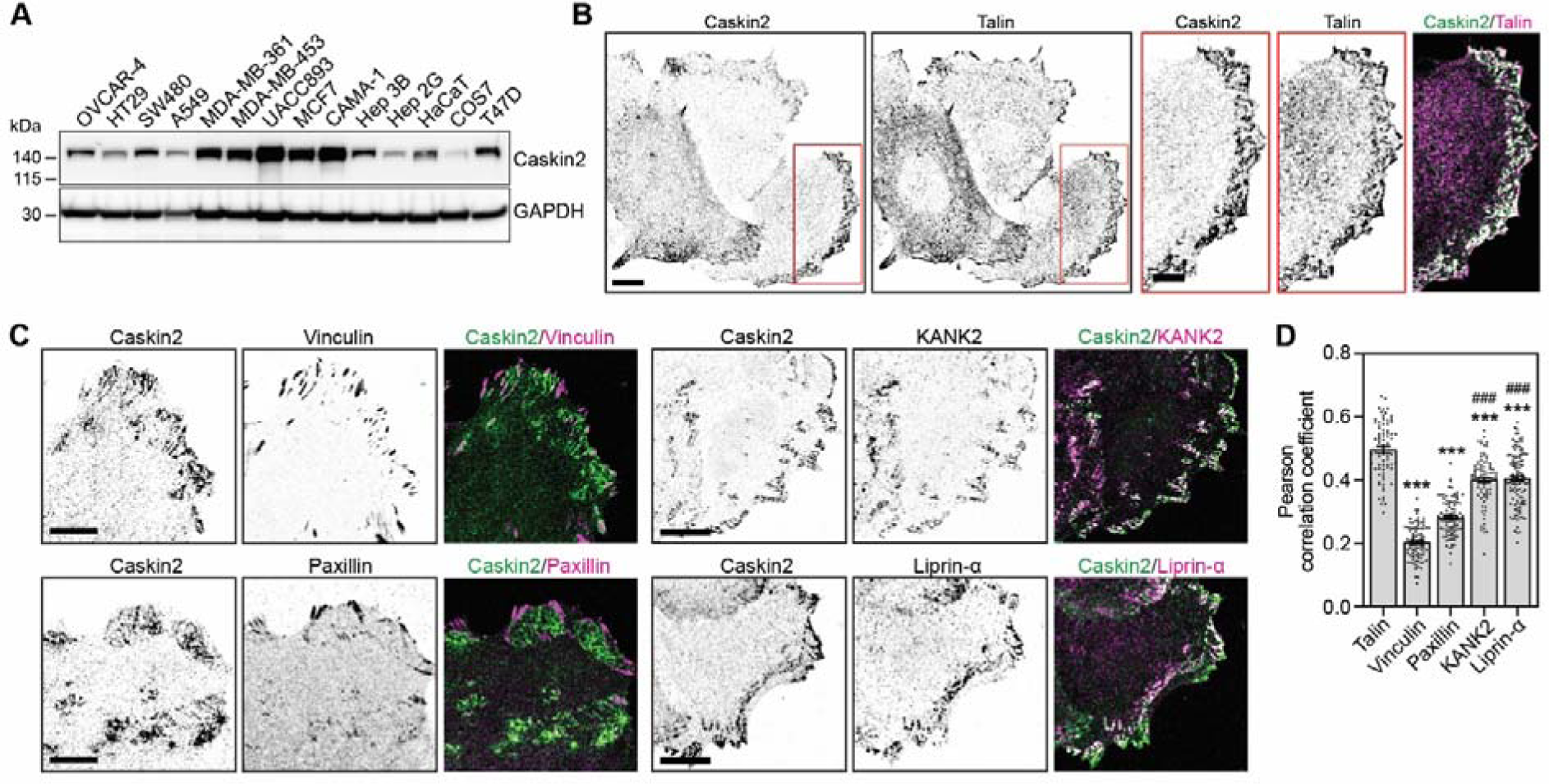
Caskin2 localisation in MCF7 breast cancer cells. (***A***) Western blot analysis of whole cell lysates from indicated cell lines, probed with antibodies against Caskin2 and GAPDH. A representative western blot is shown (n=3). (***B***) Representative IF images show endogenous Caskin2 (green in merge) and talin (magenta in merge), in MCF7 cells. Scale bar, 10 μm; in inserts represents 5 µm. Note that Caskin2 and talin do not co-localize in FAs, but co-localize in membrane-associated lattices. (***C***) Representative IF images show endogenous Caskin2 (green) together with vinculin, paxillin, KANK2 or Liprin-α (magenta) in MCF7 cells. Scale bar, 10 μm. (***D***) Pearson’s correlation analysis for the extent of co-localization of Caskin2 with indicated FA and CMSC proteins in MCF7 cells. N = 75 (Talin); 87 (Vinculin), 91 (Paxillin), 83 (KANK2), and 115 (Liprin-α) cells. *** indicates p<0.0001 compared against talin; ### indicates p<0.0001 compared against vinculin; One-way ANOVA with Holm-Šídák’s multiple comparisons test.

We examined the localization of Caskin2 in MCF7 (Fig. 2B) and UACC893 (Fig. S3A) cells using confocal microscopy. Endogenous Caskin2 localized in lattice-like structures at the cell-substratum interface, and co-staining with the FA protein talin revealed co-localization at these membrane lattices (Fig. 2B). Intriguingly, however, Caskin2 and talin did not co-localize at the ‘core’ of the FAs, with Caskin2 localized at the FA belt (Fig. 2B, Fig. S3A). In line with this finding, treatment of MCF7 cells with the myosin II inhibitor blebbistatin (10 µM) to disassemble FAs increased overall Caskin2-talin co-localization (Fig. S3B). The localization of Caskin2 to the FA belt was reminiscent of KANK1/2 localization, which is a component of cortical microtubule stabilizing complexes (CMSCs) that regulate actin-microtubule crosstalk at cell-ECM adhesion sites (Astro and de Curtis, 2015; Bouchet et al., 2016; Lansbergen et al., 2006; van der Vaart et al., 2013). Examining the co-localization of Caskin2 with other markers of FAs (vinculin and paxillin, which localize to the high-tension ‘core’ of the FA) or CMSCs (KANK2 and Liprin-α) revealed complete exclusion of Caskin2 from the FA core; meanwhile Caskin2 co-localized with KANK2 and Liprin-α, albeit to a lesser degree than with talin (Fig. 2C,D).

Tensin3, which binds directly to talin and localizes at both FAs and centrally-located, tension-independent fibrillar adhesions (Atherton et al., 2022; Clark et al., 2010), was also identified as a Caskin2-proximal protein (Fig. 1B). In MCF7 cells, Caskin2 showed little/no overlap with tensin3 at peripheral FAs, while it co-distributed with tensin3 in centrally-localized adhesions (Fig. S3C). We also observed some co-localization of integrin β5 with Caskin2 at the membrane lattices, which were distinct from integrin β5-containing flat clathrin lattices (Fig. S3D).

Based on our confocal microscopy observations we conclude that Caskin2 is excluded from the core of the FA and is localized to the FA belt as well as adjacent to the FA together with other CMSC/PMAP components, and at additional membrane patches. Therefore, Caskin2 is likely a novel component of the CMSC/PMAP and may function to regulate actin-microtubule crosstalk, similar to KANK1/2 (Bouchet et al., 2016; Sun et al., 2016).

### Caskin2 associates with Abi1 and other WAVE Regulatory Complex components

To explore the function of Caskin2 in cells, we examined the localization of Caskin2-FL-GFP in the GE11 ^tetON^ ^β1^ cells. Caskin2-FL-GFP could be observed localizing at the cell periphery but localized only weakly at integrin β1-positive adhesion structures at the cell centre (Fig. 3A). The localization at the periphery was not dependent on the talin interaction, since Caskin2-ΔLD-GFP showed a similar distribution, but lacked localization to integrin β1-positive adhesions (Fig. 3A).

**Fig. 3.**
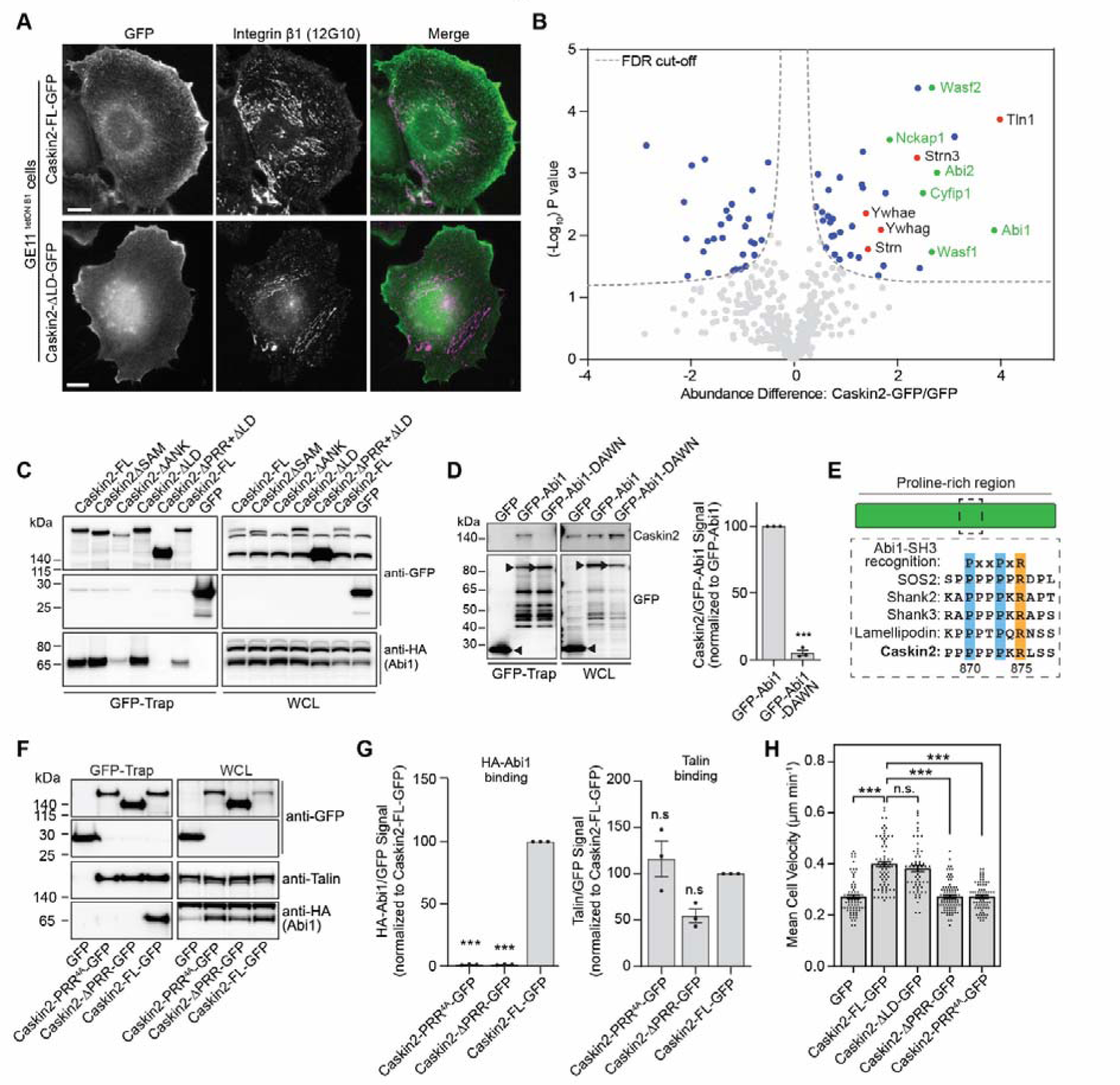
Caskin2 interacts with components of the WAVE regulatory complex (WRC) and binds directly to Abi1. (***A***) Representative IF images show Caskin2-FL-GFP and Caskin2-ΔLD-GFP expressed in GE11 ^tetON^ ^β1^ cells stained for (active) integrin β1 (12G10 antibody). (***B***) GFP-Trap pull-down of GFP (control) and GFP-tagged Caskin2 stably expressed in GE11 ^tetON^ ^β1^ cells. Pulled down proteins were identified by mass spectrometry. The volcano plot shows the results from three independent experiments (FDR: 0.05 and S0: 0.1). Significant interactors of Caskin2-GFP and GFP are indicated in blue (GFP interactors on the left side and Caskin2-GFP interactors on the right side) and red, with WAVE components indicated in green. (***C***) GFP-Trap pull-down of COS-7 cells expressing the indicated GFP-tagged Caskin2 construct, or GFP only, together with HA-Abi1. HA-Abi1 and GFP-tagged constructs were detected in the pull-down and WCL samples by Western blotting, probed with antibodies against HA and GFP, respectively. (***D***) GFP-Trap pull-down of GFP (control), GFP-tagged Abi1 and Abi1-DAWN transiently expressed in COS-7 cells. Caskin2 and GFP-tagged Abi1 and Abi1-DAWN in the GFP-Trap pull-down samples and WCLs were detected by Western blotting, probed with antibodies against Caskin2 and GFP. Black arrows indicate the expressed proteins. Error bars are standard deviation; *** indicates p<0.001 (t-test). (***E***) Cartoon showing the PxxPxR site within the PRR of Caskin2 and homology with other known binding partners of the Abi1 SH3 domain. (***F***) GFP-Trap pull-down experiments using lysates of COS-7 cells co-expressing GFP (control), Caskin2-PRR^4A^-GFP, Caskin2-ΔPRR-GFP, or Caskin2-FL-GFP together with HA-Abi1. Precipitated proteins were analyzed by Western blotting, probed with antibodies against GFP, talin and HA. (***G***) Quantification of HA-Abi1 or talin binding; error bars are standard deviation; *** indicates p<0.001, One-way ANOVA with Uncorrected Fisher’s LSD multiple comparisons test. (***H***) Quantification of cell motility of GE11 ^tetON^ ^β1^ cells. *** indicates p<0.0001, Kruskal-Wallis test with Dunn’s multiple comparisons test; n = 79 (GFP), n = 74 (Caskin2-FL-GFP), n = 61 (Caskin2-ΔLD-GFP), n = 89 (Caskin2-PRR^4A^-GFP), n = 77 (Caskin2-ΔPRR-GFP) cells, pooled from 2 independent experiments. Western blots in C, D and F are representative of 3 independent experiments.

This distribution was reminiscent of the talin-binding protein lamellipodin (Lee et al., 2009), which binds to Abi1 and other components of the WAVE Regulatory Complex (WRC) (Law et al., 2013). We explored the possibility that Caskin2 might behave similarly by using affinity purification mass spectrometry (AP-MS) to identify Caskin2-FL-GFP binding partners. In addition to Talin1, we also identified components of the WAVE complex (including Abi1/Abi2, Wasf1, Wasf2, Nckap1 and Cyfip1), several members of the 14-3-3 family of regulatory proteins (Ywhae, Ywhag and Ywhah), and members of the Striatin-interacting phosphatase and kinase (STRIPAK) family (Striatin, Striatin-3) as significant hits (Fig. 3B, Supplemental Table 3). Several of these interactors (ABI2, WASF1, WASF2, NCKAP1) were also identified in AP-MS experiments conducted in HaCaT cells stably expressing Caskin2-GFP (Fig. S4A, Supplemental Table 3).

Abi2, which is over 90% sequence identical to Abi1, has been previously shown to bind to the proline-rich region (PRR) of Caskin1 (Balazs et al., 2009). To explore the possibility that Abi1 might bind directly to Caskin2, we generated Caskin2 deletion constructs (fused to GFP at their C-terminus) and co-expressed them in COS-7 cells together with HA-Abi1. Abi1 was efficiently pulled-down by Caskin2-FL-GFP and deletion mutants containing the PRR, however, Caskin2 lacking the PRR did not pull-down HA-Abi1 (Fig. 3C). Abi1 contains an SH3 domain that commonly recognizes proline-rich sequences (Innocenti et al., 2004; Li, 2005). A GFP-tagged Abi1 mutant harboring the mutations D453A and W455N (GFP-Abi1-DAWN) that abolished binding of the Abi1 SH3 domain to proline-rich target proteins (Innocenti et al., 2004) was unable to precipitate Caskin2, unlike the WT GFP-Abi1 (Fig. 3D), demonstrating that the Abi1-Caskin2 interaction is mediated by the SH3 domain of Abi1.

Examination of the PRR of Caskin2 identified a type II peptide ligand similar to that found in other Abi1-binding partners including SOS1/2, Shank2/3, and Lamellipodin (PxxPxR; amino acids 870-875; Fig. 3E) that could potentially be recognized by the Abi1 SH3 domain. We tested this hypothesis using two different Caskin2-GFP constructs: Caskin2-GFP lacking amino acids 746-1035 (Caskin2-ΔPRR-GFP); Caskin2-GFP with PxPxxPxR mutated to AxAxxAxA (Caskin2-PRR^4A^-GFP). GFP-Trap Co-IP experiments in COS-7 cells showed that neither of these constructs could precipitate co-expressed HA-tagged Abi1, whilst talin binding was unaffected (Fig. 3F,G). Moreover, whilst Caskin2-FL-GFP co-localized with Abi1 at the cell periphery in GE11 ^tetON^ ^β1^ cells, no such co-localization was observed between Abi1 and either Caskin2-PRR^4A^-GFP or Caskin2-ΔPRR-GFP (Fig. S4B). This co-localization between Caskin2-GFP and Abi1 was not affected by the loss of talin binding, since Caskin2-ΔLD-GFP showed a similar co-localization pattern as Caskin2-FL-GFP (Fig. S4B). AP-MS experiments using these same samples (Caskin2-FL-GFP vs Caskin2-PRR^4A^-GFP transiently expressed in COS-7 cells) revealed ABI1, CYFIP1 and NCKAP1 were enriched in the Caskin2-FL-GFP compared to the Caskin2-PRR^4A^-GFP sample (Fig. S4C, Supplemental Table 4) suggesting the Caskin2-Abi1 interaction is responsible for an (indirect) association with other WRC components. We confirmed these findings by using GE11 ^tetON^ ^β1^ cells stably expressing either Caskin2-FL-GFP, Caskin2-PRR^4A^-GFP or GFP only and performing GFP-Trap co-IP experiments. Whereas Caskin2-FL-GFP precipitated endogenous Abi1, Wasf1, Cyfip1 and Nckap1, none of these were precipitated by Caskin2-PRR^4A^-GFP; 14-3-3 proteins, several of which were also detected as potential interactors in the AP-MS experiments, precipitated with both Caskin2-FL-GFP and Caskin2-PRR^4A^-GFP (Fig. S4D).

### Caskin2 promotes cell migration via its interaction with Abi1

The WAVE Regulatory Complex mediates actin polymerization downstream of Rac1 activation by activating the Arp2/3 complex, which drives the generation of branched actin filaments at cell protrusions. From our results showing that Caskin2 binds to components of the WRC, we hypothesized that Caskin2 expression may influence cell motility. We measured the migration of GE11 ^tetON^ ^β1^ cells stably expressing Caskin2-GFP, Caskin2-ΔPRR-GFP, Caskin2-PRR^4A^-GFP, Caskin2-ΔLD-GFP, or GFP only. FACS analysis confirmed the stable cells had similar levels of GFP and integrin β1 expression (Fig. S4E). Caskin2-FL-GFP expression increased cell motility compared to GFP-only expression by approximately 45% (Fig. 3H). This affect was not mediated by the Caskin2-talin interaction, since we observed no difference in motility between Caskin2-FL-GFP and Caskin2-ΔLD-GFP expressing cells (Fig. 3H). On the other hand, both Caskin2-PRR^4A^-GFP and Caskin2-ΔPRR-GFP expressing cells migrated at a similar speed as GFP-only expressing cells (Fig. 3H). Taken together, these experiments show that Abi1 recruits Caskin2 to the cell periphery, and that this interaction promotes cell migration.

### Phosphorylation of Caskin2 serine 878 regulates the interaction with Abi1

The PxxPxR SH3 domain recognition site present in Caskin2 is followed by two serine residues +2 and +3 of the arginine (S877 and S878; Fig. 3E). Intriguingly, Shank3 also has a serine +3 of the arginine in the PxxPxR site, phosphorylation of which has been shown to regulate the interaction between Abi1 and Shank3 (Perfitt et al., 2020; Wang et al., 2020a). Therefore, we wondered whether serine phosphorylation plays a similar regulatory role in the Abi1-Caskin2 interaction. To test this possibility, we generated phosphomimetic mutants of Caskin2-GFP, mutating either individual serine residues only or both serine residues together to aspartic acid residues (Caskin2-S877D-GFP, Caskin2-S878D-GFP and Caskin2-S2D-GFP, respectively). Co-IP experiments in COS-7 cells expressing these constructs together with HA-Abi1 showed HA-Abi1 binding to Caskin2-S878D-GFP was reduced by approximately 80%, whereas there was no obvious difference in binding to Caskin2-S877D-GFP (Fig. 4A,B).

**Fig. 4.**
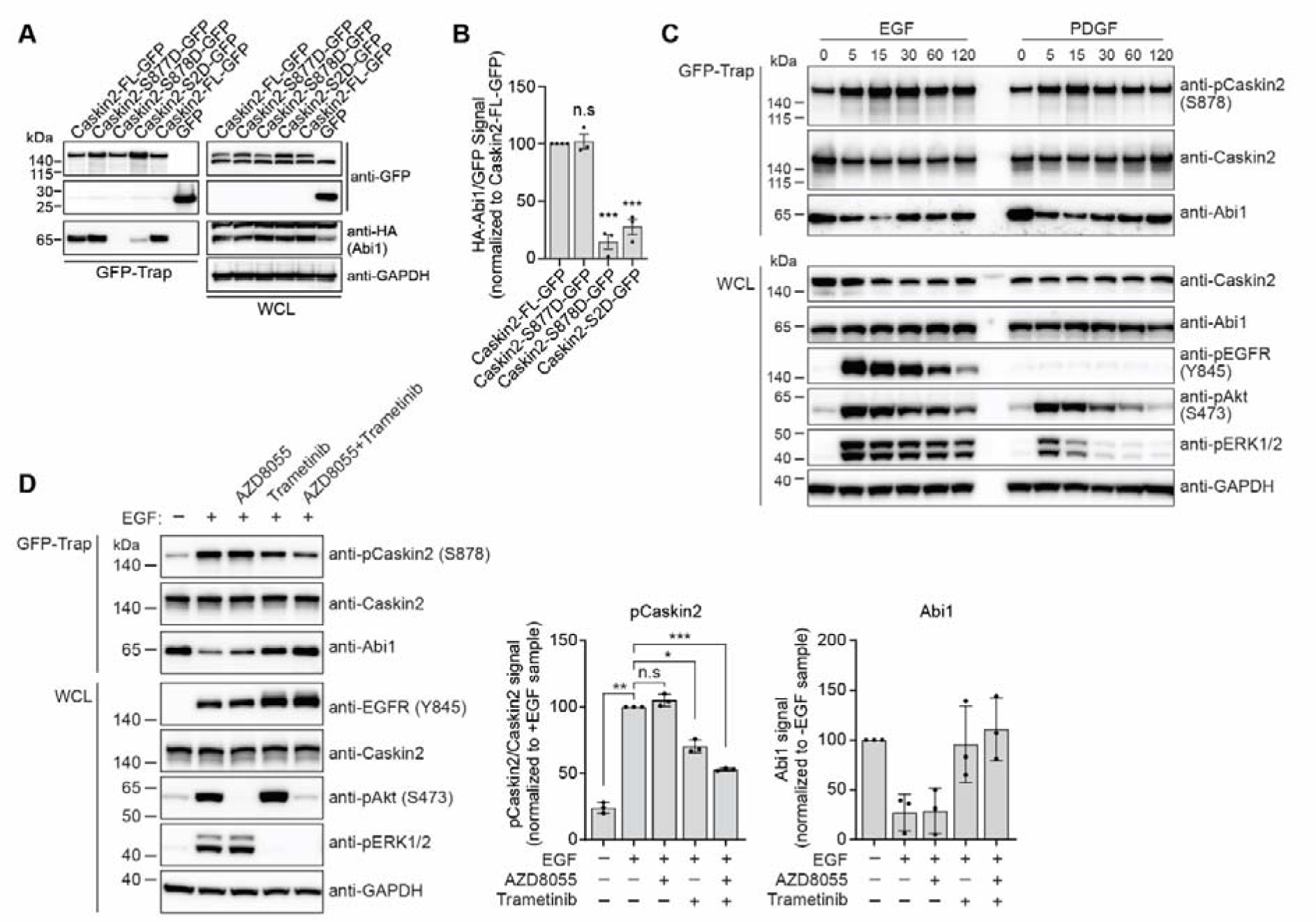
The Caskin2-Abi1 interaction is regulated by phosphorylation of S878 of Caskin2. (***A***) GFP-Trap pull-down of COS-7 cells expressing indicated GFP-tagged Caskin2 construct, or GFP only, together with HA-Abi1. HA-Abi1 and GFP-tagged constructs were detected in the pull-down and WCL samples by Western blotting, probed with antibodies against HA and GFP. (***B***) Quantification of HA-Abi1 binding. (***C***) GFP-Trap pull-down of GE11 ^tetON^ ^β1^ cells stably expressing Caskin2-FL-GFP stimulated with either EGF (100 ng/ml) or PDGF (50 ng/ml) for the indicated time. Abi1, Caskin2, and pCaskin2 (S878) were detected in the pull-down samples by Western blotting, probed with the indicated antibodies. (***D***) GFP-Trap pull-down of GE11 ^tetON^ ^β1^ cells stably expressing Caskin2-FL-GFP stimulated with EGF (100 ng/ml for 15 minutes) in the presence of AZD8055 (100 nM) or Trametinib (100 nM). Abi1, Caskin2, and pCaskin2 (S878) were detected in the pull-down samples by Western blotting, probed with the indicated antibodies. Graphs show quantification of pCaskin2 and Abi1 present in the GFP-Trap samples. Western blots in A, C and D are representative of 3 independent experiments. Error bars in B and D are standard deviation; * indicates p<0.05; ** indicates p<0.01; *** indicates p<0.001; One-way ANOVA with Holm-Šídák’s multiple comparisons test.

To explore the role of Caskin2-S878 further in regulating the Abi1 interaction, we developed a phospho-specific antibody targeting this serine. Unfortunately, high non-specific binding of the phospho-Caskin2 antibody protein sample prevented its consistent use in detecting phosphorylated Caskin2 in Western blot analysis of whole cell lysates. Only when Caskin2 was first expressed as a GFP-tagged protein and pulled down by GFP-Trap could specificity of the antibody be demonstrated. Abi1 functions downstream of growth factor receptor signaling to control Rac1-dependent actin rearrangements. Therefore, we hypothesized that Caskin2-S878 phosphorylation may occur downstream of growth factor receptor stimulation. To explore this, we conducted Co-IP experiments using GE11 ^tetON^ ^β1^ cells stably expressing Caskin2-FL-GFP stimulated with either EGF or PDGF. Both EGF and PDGF stimulation led to an increase in Caskin2 phosphorylation on S878 within 5 minutes, which was accompanied by a reduction in Caskin2-Abi1 binding (Fig. 4C). To gain insight into the kinases involved in the EGF-induced phosphorylation of Caskin2 at serine 878, we pretreated the cells with mTOR (AZD8055) and MEK1/2 (Trametinib) inhibitors for 1 h before stimulation. Both mTOR and MEK1/2 are activated downstream of EGFR activation and reduced phosphorylation of Caskin2, as detected by the phospho-specific antibody (Fig. 4D). However, only Trametinib, either alone or in combination with AZD8055 inhibited the EGF-stimulated loss of Abi1 binding. Taken together, our results indicate that Caskin2 can be phosphorylated at serine 878 by a serine/threonine kinase that acts downstream of the MEK1/2 kinases, which disrupts the interaction between Caskin2 and Abi1.

## Discussion

We have identified Caskin2 as a novel interactor of both talin and Abi1. We show that Caskin2 interacts with the R8 domain of talin via hydrophobic residues of the LD motif at the C-terminus of Caskin2; furthermore, Co-IP experiments (Fig. S1H, I) and AlphaFold modelling (Fig. S2) suggest that additional interactions can occur with the talin rod domains R2, R3, R11 and R12. Additionally, Caskin2 interacts with Abi1 via the Abi1-SH3 domain binding to the PRR of Caskin2, which connects Caskin2 indirectly to other WRC components (Fig. S4C,D). The interactions with talin and Abi1 are not mutually exclusive *in vitro*, since disrupting the Caskin2-Abi1 interaction did not affect talin binding and vice versa. (Fig. 3F,G). Mechanistically, we show that the Abi1 interaction is negatively regulated by phosphorylation of S878, which occurs downstream of MEK1/2 in response to growth factor stimulation (Fig. 4D). Intriguingly, ERK was previously shown to phosphorylate Abi1 and WAVE2, promoting the WRC-Arp2/3 interaction thus increasing cell protrusion (Mendoza et al., 2011). Therefore, ERK-mediated phosphorylation of Caskin2 may act as an additional regulatory step in this cascade by promoting the release of Abi1 from Caskin2 facilitating WRC activation and Arp2/3 binding.

Caskin2 localization was observed in two distinct compartments: i) at membrane ‘lattices’ in MCF7 cells, which contained talin and other CMSC components (Fig. 2); and ii) at the cell periphery when expressed as a GFP-fusion protein in GE11 ^tetON^ ^β1^ cells (Fig. 3). The differences in localization could arise from intrinsic differences in intracellular signaling that influence Caskin2 phosphorylation levels. Additional regulation could come from interactions of members of the 14-3-3 family of regulatory proteins and/or components of the Striatin-interacting phosphatase and kinase (STRIPAK) signaling complex, which were consistently identified in our AP-MS experiments (Supplementary Table S1, S3, S4). Expression of Caskin2-GFP was also found to increase cell motility, a phenotype that required the interaction with Abi1, but not talin (Fig. 3H). Precisely how this occurs in response to different stimuli, and whether/how Caskin2 influences actin polymerization, will require further study.

The finding that talin co-localized with Caskin2 outside of FAs in MCF7 and UACC893 cells was surprising, particularly given the absence of other FA proteins such as vinculin and paxillin. Current models suggest that vinculin binds talin when talin is in an active, open conformation (Goult et al., 2018), suggesting talin bound to Caskin2 at these sites may be in a closed conformation. Therefore, we speculate that Caskin2 may act to dampen the unfolding of talin rod domains in response to force. In this respect, Caskin2 would resemble KANK 1 and 2, which are talin-interacting proteins localizing to both CMSCs and the FA belt, which act to reduce traction forces by destabilizing the integrin-actomyosin linkage (Sun et al., 2016). The talin R2, 3, 8, 11 and 12 domains contain binding sites for vinculin and actin and binding of Caskin2 to these helical bundles may act to prevent vinculin from binding, or modulate the force required to unfold the helical bundles to expose the vinculin binding site. Force-mediated unfolding of talin rod domains may act to displace Caskin2, similar as observed for the talin binding protein RIAM (Goult et al., 2013b; Vigouroux et al., 2020).

Finally, we note that both the C-terminal LD motif and the PxxPxR Abi1-interaction motif are also present in Caskin1. Caskin1 expression is restricted to neuronal tissue where it functions as a synapse protein (Katano et al., 2018), co-localizing with Shank2/3 in the post-synaptic density (PSD) (Bencsik et al., 2019). Double knockout of Caskin1 and 2 disrupts the morphology of hippocampal dendritic spines, and impairs novel object recognition and spatial memory. Intriguingly, talin has been proposed to function in synaptic junctions by acting as a mechanical regulator of memory, with the unfolding of rod domains in response to force acting to control the association of scaffolding and signaling proteins (Barnett and Goult, 2022; Goult, 2021; Goult et al., 2021). Caskin1/2 may function to control the unfolding of talin in response to force, as discussed above, and/or may link to other signaling platforms within the synapse (such as the scaffolding protein CASK for Caskin1 (Smirnova et al., 2016)).

## Acknowledgements

We thank Lodewyk Wessels (Netherlands Cancer Inst., Amsterdam, the Netherlands), Sylvia Le Dévédec (Leiden University, Leiden, the Netherlands), Christoph Ballestrem (Wellcome Trust Centre for Cell-Matrix Research, University of Manchester, Manchester, UK) and Metello Innocenti (Heidelberg University Biochemistry Center (BZH), Heidelberg University, Germany) for sharing cell lines and reagents, and Stefan Uebel (Max Planck Institute fo Biochemistry, Martinsried, Germany) for help with the biophysical analyses. This work was supported by the Dutch Cancer Society (project number 12143) and the Netherlands Organization for Scientific Research (NWO) as part of the National Roadmap Large-scale Research Facilities of the Netherlands, Protein@Work (project number 184.032.201). A. Sonnenberg and R. Fässler would like to thank the Alexander von Humboldt Foundation for supporting A. Sonnenberg during his sabbatical year at the Max Planck Institute.

## Authors contributions

W. Wang, P. Atherton, and A. Sonnenberg conceived the study, designed experiments, interpreted data, and wrote the manuscript with contributions from A. Perrakis. W. Wang, P. Atherton, M. Kreft, L. te Molder, L. Hoekman, P. Celie, R. P. Joosten, and A. Sonnenberg performed and analyzed the experiments. R. Fässler provided resources and reviewed and edited the manuscript.

## Materials and Methods

### Construction of expression plasmids

Human Caskin2 cDNA was cloned into the *Not*I and *Sal*I sites of a pCMV-SPORT6 vector (Horizon, MHS6278_-_202759048). Caskin2 cDNA (with the stop codon removed) was assembled by *in vitro* ligation of an N-terminal part of 3.7 kb and a C-terminal PCR fragment, and cloned into the pUC19-HA-SKG269-BirA* and pUC19-HA-SKG269-GFP plasmids (Zuidema et al., 2022b), replacing the HA-tagged SKG269 DNA. Deletion mutants of Caskin2 and Caskin2-4A were generated by site-directed mutagenesis with the PCR-based overlap extension method using Pfu DNA polymerase (Promega), and fragments containing the different mutations were exchanged with corresponding fragments in the pUC19-CASKIN2-GFP plasmid. Caskin2-S877, Caskin2-S877/878D (S2D) and Caskin2-PRR^4A^ constructs were generated by purchasing gene blocks containing the desired point mutations (Genewiz) flanked with HindIII and NheI restriction sites which were used to clone the fragments into Caskin2-FL. Retroviral vectors containing mutant Caskin2-GFP and Caskin2-ΔLD cDNAs were generated by subcloning the mutant Caskin2 cDNA into the *EcoR*I and *Not*I restriction sites of LZRS-MS-IRES-Zeo (Kinsella and Nolan, 1996). Wild-type Caskin2 and Caskin2-4A were cloned into the mammalian expression vector pcDNA3 using the *EcoR*I restriction sites.

### Cell culture and generation of cell lines

Immortalized PA-JEB keratinocytes stably expressing integrin β4 (PA-JEB/β4) were generated by retroviral transduction, as previously reported (Sterk et al., 2000). Cells were maintained in keratinocyte growth medium (KGM; Invitrogen) supplemented with bovine pituitary gland extract (50 μg ml^-1^), EGF (5 ng ml^-1^), and streptomycin/penicillin (100 units ml^-1^; Sigma-Aldrich). One day prior to beginning the experiment, the medium was changed into Dulbecco’s modified Eagle’s medium (DMEM; Gibco) containing 10% fetal bovine serum (FBS; Serana Europe GmbH, Pessin, Germany) and 100 units ml^-1^ each of streptomycin and penicillin. GE11 ^tetON^ ^β1^ were generated by infecting GE11 cells (Gimond et al., 1999) with a retroviral vector expressing integrin β1 under a tet-inducible promotor (Retro-X™ Tet-On®; Clontech, USA). Integrin β1-induced and FACS sorted cells were cultured in DMEM with 10% fetal bovine serum, containing 1 μg/ml doxycline. Human breast cancer cell line MCF7 (a gift from Dr. L. F. A. Wessels, the Netherlands Cancer Institute, the Netherlands), UACC893 (a gift from Dr. S. E. Le Dévédec, Leiden University, the Netherlands) and COS-7 cells were cultured in DMEM/FCS. All cells were maintained at 37°C in a humidified atmosphere of 5% CO_2_. All cell lines were routinely tested for mycoplasma contamination every three months.

Stable cell lines stably expressing GFP-tagged wild-type and mutant Caskin2 constructs were generated by retroviral infection. Retroviral expression plasmids were transfected into Phoenix packaging cells using the calcium phosphate precipitation method and virus containing supernatant was collected 72 h post-transfection (Kinsella and Nolan, 1996). After infection overnight at 37°C, infected cells were selected with 0.2 mg/ml zeocin (Invitrogen) for two weeks (Wang et al., 2020b). GFP positive cells were isolated by fluorescence-activated cell sorting (Beckman Coulter Moflo Astrios cell sorter) for following use.

For transient expression, COS-7 cells were grown to 70% confluence in 10 cm culture dishes and transfected with plasmid DNAs using the DEAE-dextran method (Schaapveld et al., 1998). Briefly, 4.5 µg of plasmid DNA was mixed with 500 µg/ml DEAE-dextran in 2.5 ml PBS. Cells were washed two times with PBS and then incubated for 30 min at 37°C with the DEAE-dextran solution. Subsequently, 80 µM chloroquine was added to the cells in 10 ml DMEM/FCS and cells were incubated at 37°C for 3 h. After aspiration of the medium, the cells were treated with 3 ml of DMEM/FCS containing 10% DMSO for 2.5 min. The DMSO solution was then removed and replaced with fresh medium, after which the cells were further incubated for 48-72 h before analysis.

### Antibodies used for immunofluorescence, western blotting and FACS

Primary antibodies used are listed in Table S5. The affinity-purified rabbit antibody selective for Caskin2 (pS787) was generated by Davids Biotechnologie GmbH (Regensburg, Germany), using the phospho-peptide PPPKRLSS(P)VSGPSPEPPPLD as immunogen. The phospho-peptide was synthesized by the protein core facility of the Max-Planck Institute (Martinsried, Germany). Rabbits were immunized with the phospho-peptide coupled to Keyhole Limpet Hemocyanon (KLH) using m-maleimidobenzoyl-N-hydroxysuccinimide (MBS). Affinity chromatography was performed using a phospho-peptide matrix. Antibodies reactive with the non-phospho version of the protein antigen were removed by passing the affinity purified antibodies over an affinity matrix with non-phospho-peptide.

Secondary antibodies for IF were: goat anti-rabbit Alexa Fluor 488, goat anti-mouse Alexa Fluor 561, goat anti-rat Texas Red, goat anti-mouse Alexa Fluor 647 (Invitrogen; 1:200). Secondary antibodies for Westering blotting were goat anti-mouse IgG HRP (Bio-Rad; 1:3000) and polyclonal goat anti-rabbit IgG HRP (Bio-Rad; 1:5000). Secondary antibodies for FACS were goat anti-mouse Alexa Fluor 647 (Invitrogen; 1:200).

### Immunofluorescence and image analysis

PA-JEB/β4 keratinocytes were seeded on glass coverslips and cultured for 24 h in complete KGM, and then continued to be cultured for a further 16 h in DMEM with 10% FCS. GE11 ^tetON^ ^β1^ and MCF7 cells were seeded on glass coverslips and cultured for 24 h in DMEM with 10% FCS. Cells were fixed in 2% paraformaldehyde and permeabilized with 0.2% Triton X-100 for 5 min, blocked with PBS containing 2% bovine serum albumin (BSA; Sigma-Aldrich) for 45 min and subsequently incubated with primary and secondary antibodies for 1 h. In between antibody steps, the coverslips were washed three times with PBS. Actin fibers were visualized by staining with Phalloidin Alexa Fluor 647 (Cell Signalling Technology). Nuclei were stained with 4, 6-diamidino-2-phenylindole (DAPI). Samples were mounted on glass slides in Mowiol after three washing steps with PBS. Images were acquired with a Leica TCS SP5 confocal microscope with a 63x (NA 1.4) oil objective. Software for image acquisition is Leica LAS AF 3.0.2 (Wang et al., 2020b).

Imaging analysis was performed using FIJI (Schindelin et al., 2012). Co-localization efficiency of Caskin2 and various proteins, including integrin β4, focal adhesion proteins and CMSC components, was performed using Pearson’s co-efficient analysis in the JACoP module of ImageJ (Carisey et al., 2013; Wang et al., 2020b).

### Western blotting, immunoprecipitations and GFP-Trap assay

For the analysis of phospho-MLC levels in whole cell lysates, cells were grown to 70-80% confluence and lysed in RIPA buffer, supplemented with Na_3_VO_4_ (1.5 mM), NaF (15 mM), protease inhibitor cocktail (1:1000; Sigma-Aldrich) and Phosphatase Inhibitor Cocktail 3 (1:100; Sigma-Aldrich). Lysates were clarified by centrifugation at 14,000 x *g* for 1h at 4°C, and then mixed at a ratio of 1:1 with 4x SDS sample buffer (200 mM Tris–HCl pH 6.8, 8% SDS, 40% glycerol, 50 mM EDTA, 0.4% Bromophenol Blue, 2% β-mercaptoethanol) and then heated at 95 °C for 5 min. A small portion of lysate was reserved for quantification of protein concentration using the Pierce™ BCA Protein Assay kit, following the manufacturer’s instructions. Proteins were resolved by sodium dodecyl sulfate polyacrylamide gel electrophoreses (SDS-PAGE) on 15% homemade gel and transferred to Immobilon-P transfer membranes (Millipore Corp).

Cell lysates, prepared by using Nonidet P-40 lysis buffer (20 mM Tris HCl pH 7.5, 100 mM NaCl, 1% NP-40) with protease and phosphatase inhibitors were incubated with GFP-Trap Agarose beads (Chromotek) in a rotation wheel for 6 h at 4°C. Bound proteins and input samples (whole cell lysates) were loaded on Novex NuPAGE 4-12% gradient Bis-Tris gel (Invitrogen) and transferred to Immobilon-P transfer membranes. Membranes were blocked for at least 2 h in 2% BSA in TBST (10 mM Tris (pH 7.5), 150 mM NaCl, 0.05% Tween 20) before incubation with primary antibody overnight at 4°C and with secondary antibody for 1 h at room temperature. After each incubation step, the membranes were washed twice with TBST and twice with TBS (TBST without Tween 20). Finally, proteins were visualized using Clarity™ Western ECL Substrate (Bio-Rad Laboratories, Inc.), as described previously (Wang et al., 2020b).

### BioID assay

PA-JEB/β4 cells stably expressing Caskin2-BirA* fusion proteins were grown to 70%-80% confluence on 3 × 145 mm plates in biotin-depleted keratinocyte growth medium before they were treated with 50 μM biotin (Sigma-Aldrich #B4501) for 20 h at 37°C in DMEM/10% FCS. Biotinylated proteins were captured with Streptavidin Sepharose beads (GE Healthcare) and bound proteins analyzed by mass spectrometry.

### MST measurements

MST measurements were carried out using premium-coated capillaries to reduce non-specific interaction of the proteins with the glass surface on a Monolith NT.115 red-blue (Nanotemper, Munich, Germany), as described previously (Zuidema et al., 2022b). Peptides and proteins were transferred into MST buffer (20 mM Tris, pH 7.5, 200 mM sodium chloride, 1 mM TCEP, 0.05% Tween-20) to avoid artifacts derived from buffer mismatches. Atto488-labeled Caskin2-LD and scrambled peptide (synthesized by MPIB Core Facility) were used as ligands at a concentration of 50-200 nM and added in a 1:1 ratio to a 1:1 serial dilution of recombinant R7R8. The measurements were performed at 10 to 20% LED power and 20 and 40% MST power, and finally analyzed using the MO Affinity Analysis Software (Nanotemper).

### Recombinant protein expression and purification

A codon-optimized synthetic gene encoding murine talin-R7/R8 domains (residues 1358-1653) was cloned into the pETNKI-6xhis-3C-LIC vector (Luna-Vargas et al., 2011). The 6xhis-3C-talin-R7/R8 protein was expressed in Bl21(DE3) cells, for 18 hours at 20 °C upon induction with 0.4 mM IPTG. Cells were harvested by centrifugation and stored at -20 °C. After thawing, cells were resuspended in lysis buffer (25 mM Tris (pH 8.0), 10 mM imidazole, 200 mM NaCl, 1 mM TCEP, 5 μg/mL Dnase) and lysed by sonication. The soluble lysate fractions were collected after centrifugation (55,000g at 4 °C for 30 minutes) and applied to a Nickel Sepharose column. Beads were washed with wash buffer (lysis buffer without DNAse) and protein was eluted in the same buffer supplemented with 250 mM imidazole. The 6xHis tag was cleaved off by his-3C protease during dialysis against 25 mM TRIS pH 8.0,100 mM NaCl, 1 mM TCEP for 16 hours at 4 °C. The protein solution was diluted with an equal volume of 25 mM Tris pH 8.0, 1 mM TCEP before his-3C protease and uncleaved 6xhis-Talin-R7/R8 were removed by Nickel Sepharose. The protein was further purified by anion-exchange chromatography (Resource Q column, Cytiva) followed by size exclusion chromatography (S75 Superdex 16/60, Cytiva) in 25 mM TRIS (pH 8.0), 100 mM NaCl, 1 mM TCEP. Talin-R7/R8 eluted in a single peak. Protein was concentrated to 32 mg/ml and aliquots were snap-frozen in liquid nitrogen and stored at -80 °C.

### Talin-R7/R8 – Caskin2-LD peptide crystallization

Purified talin-R7/R8 protein (32 mg ml^-1^) was mixed with solubilized Caskin-2 LD peptide in a 1:2 molar ratio of protein:peptide. Crystallization screening was performed using the vapor diffusion sitting-drop method; droplets of 200 nanoliters, composed of equal volumes of protein complex and crystallization agent, were prepared using a Mosquito dropsetter (SPT Labtech) as described previously (Newman et al., 2005). Conditions revealing initial crystal growth were further optimised using a Formulator liquid handler (Formulatrix). Diffracting crystals were grown in 0.1 M bis-Tris propane pH 7.0, 0.1 M NaBr and 22 - 26% PEG3350. Crystals were cryo-protected by increasing the PEG3350 concentration to 30% and by addition of 10 % glycerol before vitrifying them in liquid nitrogen.

### X-ray crystallography structure analysis

Crystallographic data were collected at the PX1 beamline at the Swiss Light Source (SLS), Villigen, Switzerland and processed with XDS (Kabsch, 2010) and Aimless (Evans and Murshudov, 2013). The phase problem was solved by molecular replacement using Phaser (McCoy et al., 2007) and two separate search models, using the Talin R7 and R8 domains (PDB: 5fzt) (Zacharchenko et al., 2016). After a few cycles of model building in Coot (Emsley et al., 2010) and model refinement in REFMAC5 (Kovalevskiy et al., 2018), the electron density for Caskin-2 peptide was clearly visible. Due to the lower local resolution of the map at that area, five probable alternative registers were modeled and refined. Additionally, the register was sought independently with the “Assign sequence” function in Coot. Both approaches led to the same register. The structure model was finalized by performing alternating cycles of model building in Coot, validation in MolProbity (Chen et al., 2010) and Tortoize (van Beusekom et al., 2018), and refinement in PDB-REDO using homology restraints (van Beusekom et al., 2018). Data collection and model statistics are reported in Table S2.

### Modelling of the interaction of the Caskin2 LD peptide in complex with talin rod domains

The ColabFold interface to AlphaFold (Mirdita et al., 2022) and AlphaFold Multimer (Evans et al., 2022; Jumper et al., 2021) were used to model the interactions of the Caskin2 LD peptide with all thirteen complexes with the talin rod domains. The sequence of the peptide was used as “chain A” and that of each rod domain as “chain B”. An artificial sequence for R7 was constructed by fusing the two sequence fragments that make up those helical fragments, based on our crystal structure. The confidence of the interactions was judged by manual inspection of the Predicted Alignment Error (PAE) plots, which is a useful metric to assess how confident the model is about the interface. The modeling procedure could reproduce our crystal structure of the R8 domain with very high confidence in all five top-ranked models, while it only produced very low confidence interfaces with the R7 domain (which does not bind the LD peptide in our structure), indicating that this is a suitable protocol to find additional interactors. High confidence complex interfaces were predicted for all five top-ranked models for the complexes with R3, R11 and R12, while two high-confidence models were available for the R2 complex.

### Mass spectrometry

As described in our previous work (Wang et al., 2020b), samples from BioID, GFP-Trap and peptide pull-down assay were separated on a 4-12% SDS-PAGE gel. The gel was stained with Coomassie Blue and lanes were excised and then reduced by treating with dithiothreitol and alkylated with iodoacetamide. After digestion with trypsin (mass spec grade, Promega), peptides were extracted with acetonitrile. A vacuum centrifuge was used to dry the digests, which were re-suspended in 10% formic acid. Peptides were analyzed by nanoLC-MS/MS on an Orbitrap Fusion Tribrid mass spectrometer equipped with a Proxeon nLC1000 system (Thermo Scientific). Subsequently, samples were eluted from the analytical column at a constant flow of 250 nl min-1 in a 140-min gradient, containing a 124-min linear increase from 6% to 30% solvent A (0.1% formic acid/water), followed by a 16 min wash at 100% solvent B (0.1% formic acid/80% acetonitrile).

For data analysis, we used either the human Swissprot database (20,432 entries, release 2019_09) or the mus musculus Swissprot database (17,141 entries, release 2023_03), as data sources and applied MaxQuant (version 2.4.2.0) (Cox et al., 2014) with standard settings to our raw data for label-free quantitation (LFQ). MS/MS data were concatenated with the reversed version of all sequences from the database. Trypsin/P was chosen as cleavage specificity allowing two missed cleavages; Carbamidomethylation (C) was set as a fixed modification, while oxidation (M) was used as variable modification. LFQ intensities were Log2-transformed in Perseus (version 1.6.7.0) (Tyanova et al., 2016), after which proteins were filtered for at least two valid values (out of 3 total). Missing values were replaced by imputation based a normal distribution using a width of 0.3 and a downshift of 1.8. Differentially expressed proteins were determined using a t-test.

### Cell Migration

Cell motility assays were conducted using time-lapse recordings of cells cultured on plastic 12-well plates in DMEM containing 1% FCS and 1% antibiotics. Phase-contrast images were acquired using a Zeiss AxioObserver Z.1 microscope equipped with a heated stage (37°C and 5% CO_2_), with a 10x/0.30 EC Plan-Neofluar Ph1 objective (Zeiss), captured with a Zeiss AxioCam MRm. Images were acquired every 10 minutes for 20 hours. Cells were tracked manually using ImageJ; tracking data was analyzed using the Chemotaxis Tool plugin (Ibidi).

### Statistical analysis

The two-sided Student T test was used to calculate significance between two groups using GraphPad Prism 10 (La Jolla). Data distribution was tested using the D’Agostino-Pearson normality test with a significance levels of 0.05. Graphs were made in GraphPad Prism; statistically significant values are indicated as *, p < 0.05; **, p < 0.01; ***, p < 0.001; ****, p < 0.0001.

### Online Supplemental Material

Fig. S1: Related to Fig. 1. Sequence alignment of Caskin2 showing the LD-motif; characterization of the Caskin2-LD interaction with talin R8.

Fig. S2: Related to Fig. 1. AlphaFold modelling of potential interactions between the Caskin2 LD-motif and additional talin rod domains.

Fig. S3: Related to Fig. 2. Localization of Caskin2 in UACC893 and MCF7 cells.

Fig. S4: Related to Fig. 3. GFP-Trap AP-MS data from experiments in HaCaT cells; data showing Caskin2-GFP interacts with several WRC proteins; FACS profiling of GFP and integrin β1 expression.

Table S1: Related to Fig. 1. Caskin2 proximal proteins identified by BioID in PA-JEB/β4 keratinocytes.

Table S2: Related to Fig. 2. Data collection and refinement statistics of the Talin R7R8/Caskin2-LD complex.

Table S3: Related to Fig. 3 and Fig. S4. Mass spectrometry analysis of Caskin2-GFP binding partners in GE11 ^tetON^ ^β1^ or HaCaT cells.

Table S4: Related to Fig. S4. Mass spectrometry analysis of Caskin2-GFP and Caskin2-PRR^4A^-GFP binding partners in COS-7 cells.

Table S5: List of primary antibodies used.

---

**Supplemental Fig. 1.**
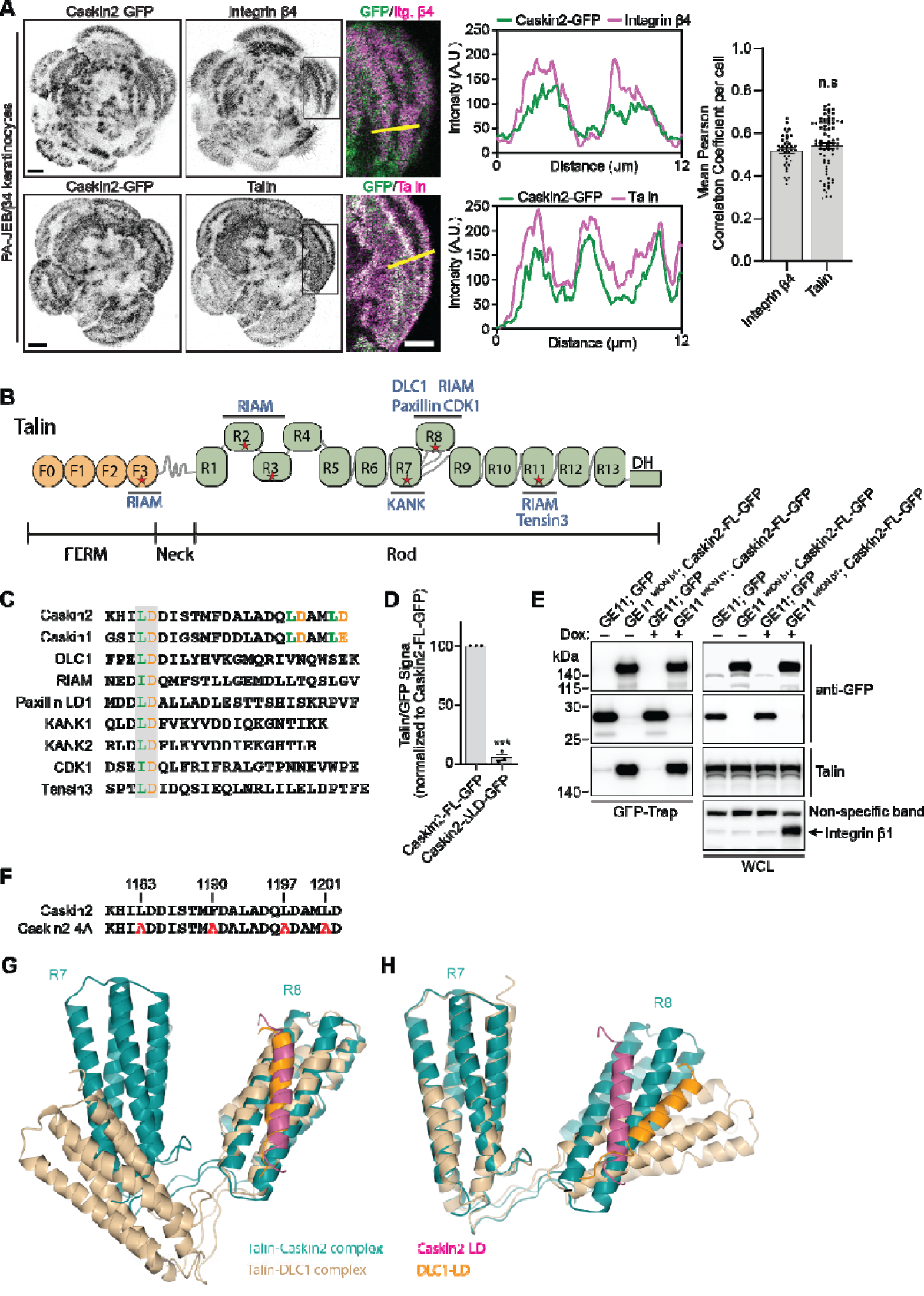
Caskin2 co-distributes with talin and localizes in the proximity of HDs, FAs and CMSCs. (***A***) Representative immunofluorescence images show Caskin2-GFP (green) together with integrin β4 or talin (magenta) in PA-JEB/β4 keratinocytes. Scale bar, 10 μm; in inserts represents 5 µm. Line profiles shows Caskin2 co-localises with both talin and integrin β4. Graph shows the mean Pearson Correlation Coefficient per cell between the Caskin2-GFP signal and either Integrin β4 or talin; n.s indicates no significant difference; error bars are standard error; unpaired t-test; n = 40 (Integrin β4) and 71 cells (talin) pooled from 3 independent experiments. (***B***) Talin comprises of a FERM domain with four subdomains (F0-F3, in orange) and a rod domain with 13 bundles (R1-R13, in green), followed by a C-terminal dimerization helix (DH). Some critical interacting proteins are indicated. (***C***) Sequence alignment of the talin-binding motifs of Caskin2, Caskin2, DLC1, RIAM, Paxillin LD1, KANK1, KANK2 and CDK1. Acidic (orange) and hydrophobic (green) residues that mediate the interaction are highlighted. (***D***) Quantification of Talin binding to the indicated GFP-tagged construct, normalized to Caskin2-FL-GFP. Values were quantified from 3 independent Western blots, error bars are standard error, *** indicates p<0.0001 (unpaired t-test). (***E***) GFP-Trap pull-down of GFP expressed in wild-type GE11 (β1-null) or Caskin2-FL-GFP expressed in GE11 ^tetON^ ^β1^ cells. Cells were cultured continually with or without doxycycline as indicated. Proteins in the GFP-Trap sample and WCL (input) are detected by Western blotting, using antibodies against GFP, talin, and integrin β1. Note that doxycycline induces integrin β1 expression only in the GE11 ^tetON^ ^β1^ cells. Talin binding to Caskin2-FL-GFP is not affected by the presence of integrin β1. A representative Western blot is shown (n=3). (***F***) Amino acid sequences of the Caskin2 LD-motif and the 4A mutant containing 4 alanine mutations. (***G*** *and **H***) Comparison of the crystal structure of talin R7R8 in complex with Caskin2-LD (magenta) and in complex with DLC1-LD (residues 467-489; orange), which is aligned on the R8 domain (***G***) and R7 domain (***H***).

**Supplemental Fig. 2.**
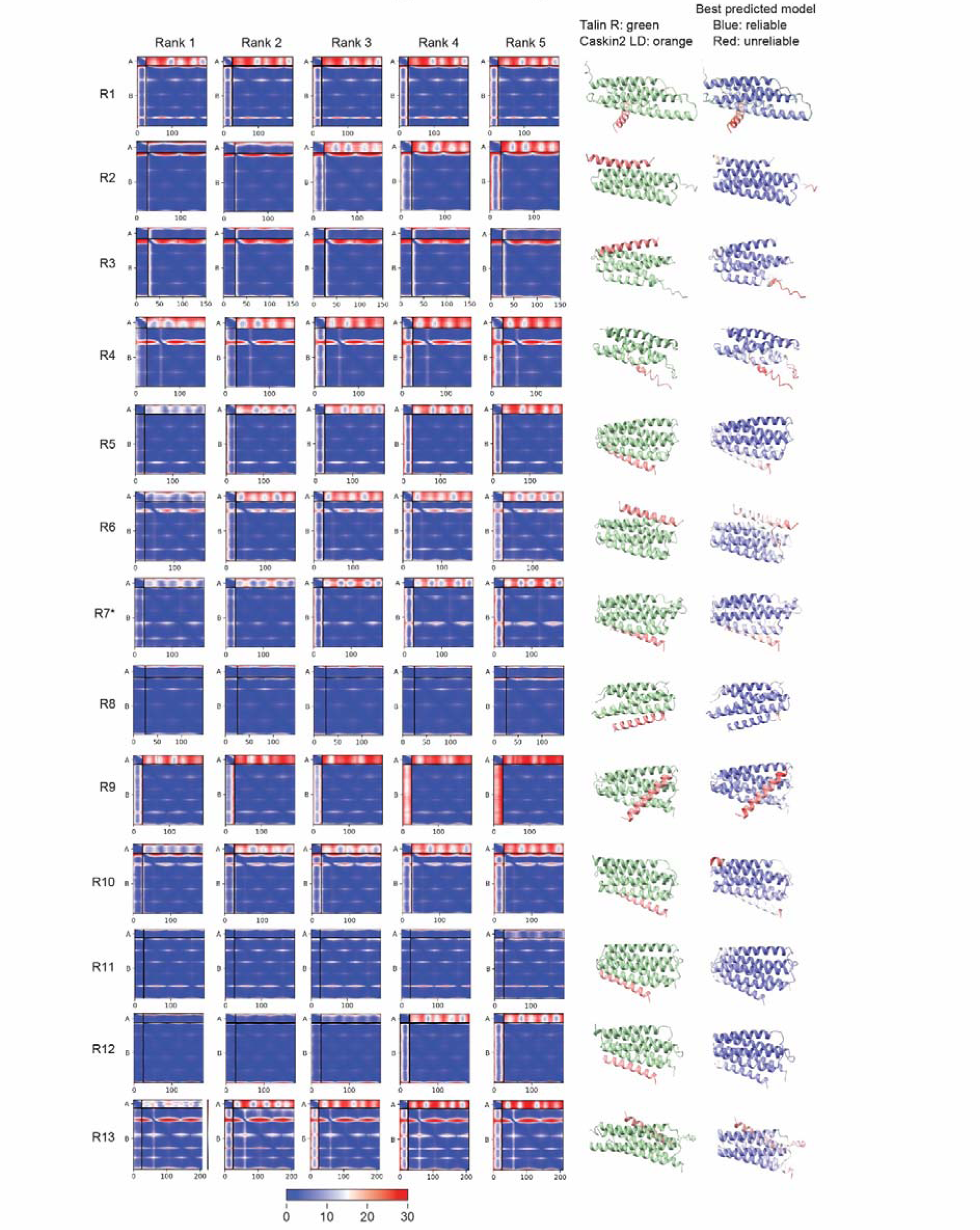
AlphaFold modelling for Caskin2-LD interactions with different talin rod domains. The different columns show: the talin rod domain examined in each row; the Predicted Alignment Error (PAE) plots for the five top-ranked models (blue: high confidence interaction; red: low confidence interaction); a cartoon model of the complex with talin rod domain in green and the Caskin2 LD peptide in orange; the same model coloured by local confidence.

**Supplemental Fig. 3.**
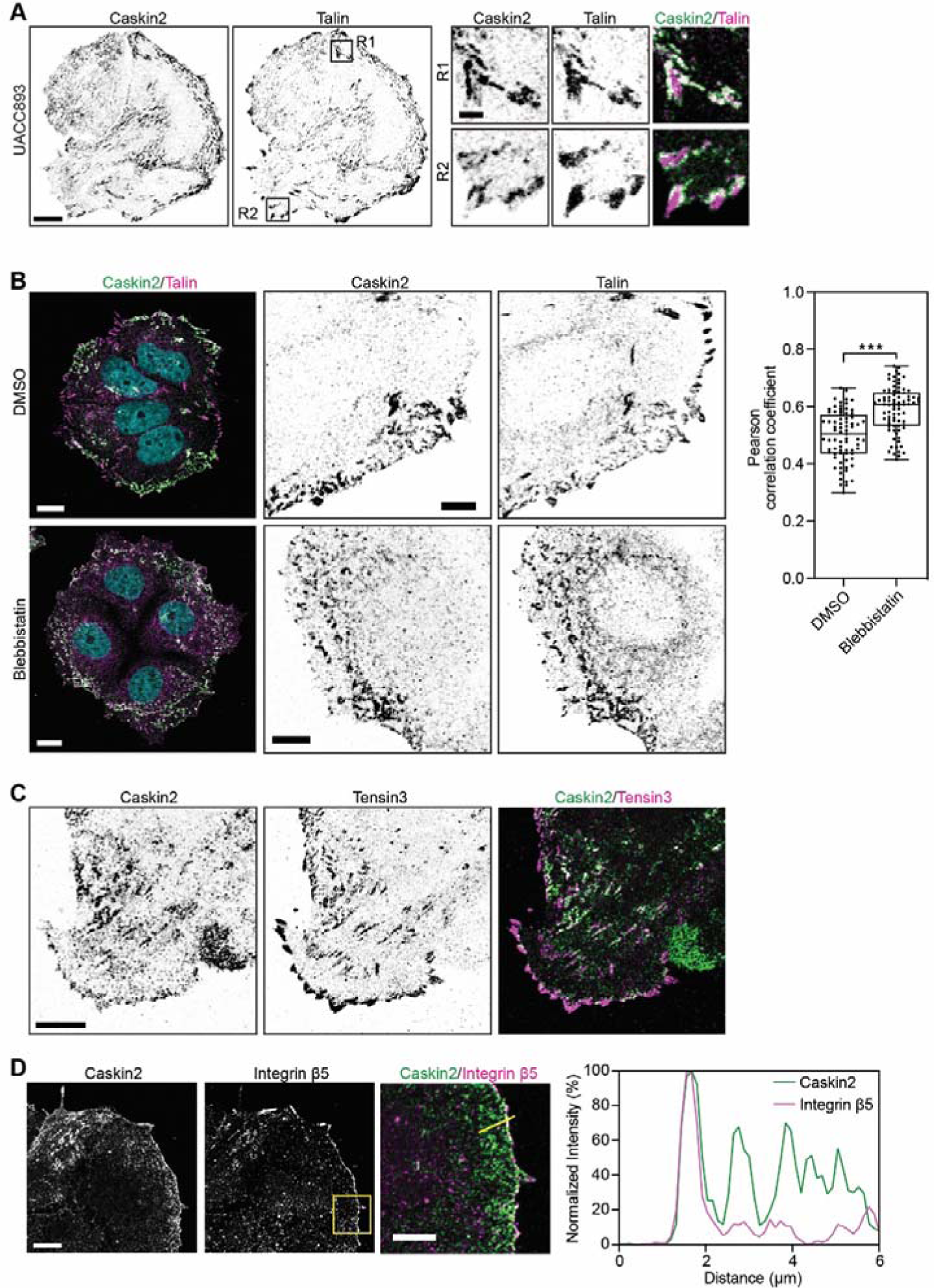
Localization of endogenous Caskin2 in breast cancer cells. (***A***) Representative IF images show endogenous Caskin2 (green in merge) and talin (magenta in merge), in UACC893 cells. Scale bar, 10 μm; in inserts represents 2 µm. (***B***) Representative IF images show Caskin2 (green), talin (magenta), and DAPI (cyan) in MCF7 cells treated with DMSO or 20 µM blebbistatin for 45 min. Scale bar, 10 μm; in magnified region, 5 µm. Pearson’s correlation analysis for the extent of co-localization of Caskin2 with talin in DMSO (n = 75) or Blebbistatin (n = 81) treated MCF7 cells. Data are presented as box-and-whisker plots, in which the box extends the 25^th^ to 75^th^ percentiles, the middle line indicates the median, and whiskers go down to the smallest value and up to the largest. ****, P < 0.0001, t-test. (***C***) Representative IF images showing endogenous Caskin2 (green) together with Tensin3 (magenta) in MCF7 cells. Scale bar, 10 μm. (***D***) Representative IF images showing endogenous Caskin2 (green in merge) and integrin β5 (magenta in merge) in MCF7 cells. Yellow arrow shows integrin β5 in flat clathrin lattices, which do not contain Caskin2. Scale bar indicates 10 µm.

**Supplemental Fig. 4.**
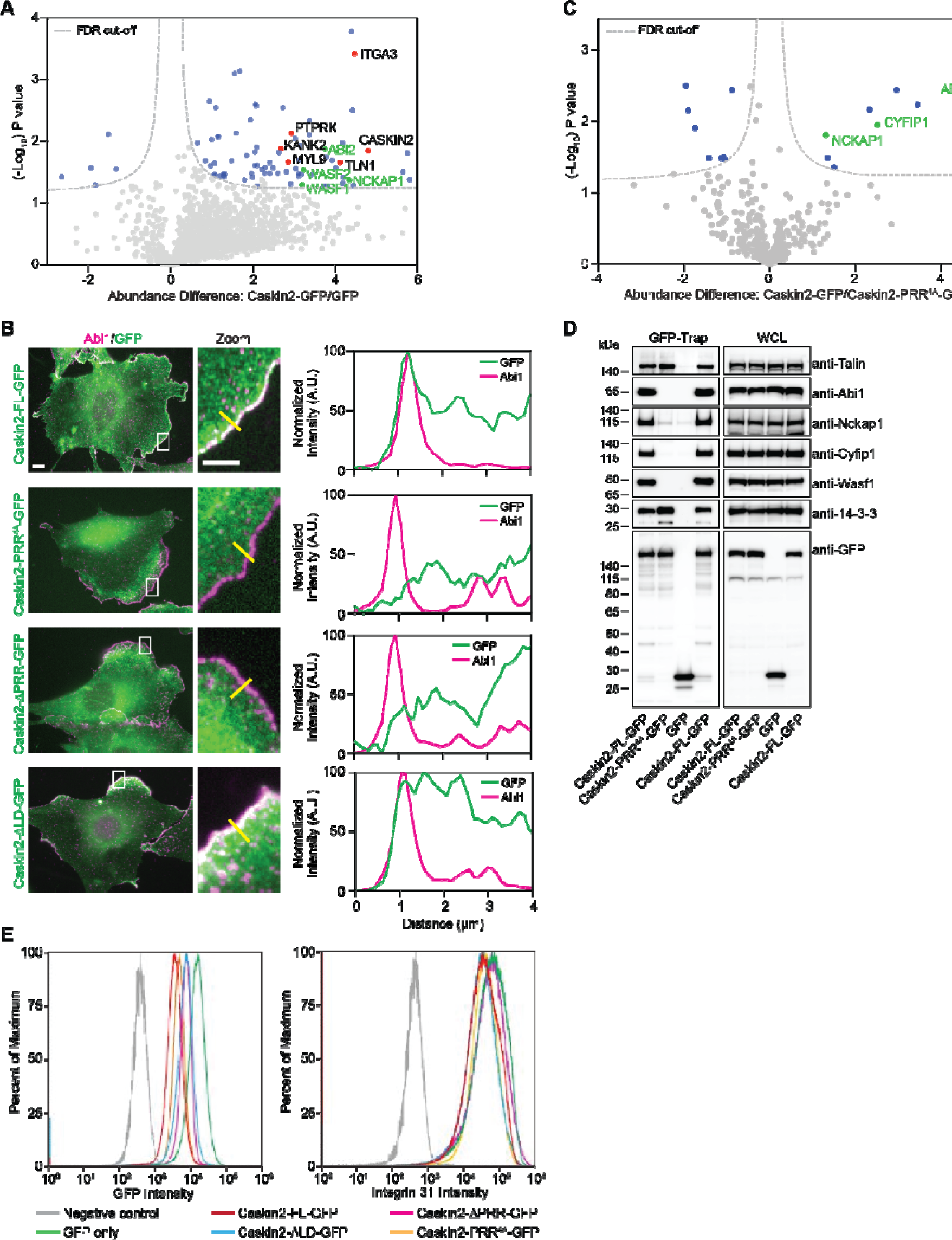
Caskin2 associates with components of the WAVE Regulatory Complex. (***A***) GFP-Trap pull-down of GFP (control) and GFP-tagged Caskin2 stably expressed in HaCaT cells. Pulled down proteins were identified by mass spectrometry. The volcano plot shows the results from three independent experiments (FDR: 0.05 and S0: 0.1). Significant interactors of Caskin2 and GFP are indicated in blue (GFP interactors on the left side and Caskin2 interactors on the right side) and red, with WAVE components indicated in green. (***B***) Representative immunofluorescence images of GE11 ^tetON^ ^β1^ cells expressing Caskin2-FL-GFP, Caskin2-PRR^4A^-GFP, Caskin2-ΔPRR-GFP, or Caskin2-ΔLD-GFP together with endogenous Abi1. Line profiles show Caskin2-FL-GFP co-localises with Abi1 at the cell periphery, which is not seen in the two constructs deficient for Abi1 binding. Deletion of the talin-binding LD motif does not affect Abi1 co-localization. Scale bars indicate 10 µm; in magnified regions indicate 5 µm. (***C***) GFP-Trap pull-down of Caskin2-FL-GFP and Caskin2-PRR^4A^-GFP co-expressed with HA-Abi1 in COS-7 cells. Pulled down proteins were identified by mass spectrometry. The volcano plot shows the results from three independent experiments (FDR: 0.05 and S0: 0.1). Significant interactors of Caskin2-FL-GFP but not Caskin2-PRR4A-GFP are on the right side indicated in blue; WAVE components are indicated in green. (***D***) GFP-Trap pull-down of GFP (control), Caskin2-FL-GFP or Caskin2-PRR^4A^-GFP stably expressed in GE11 ^tetON^ ^β1^ cells. Precipitated proteins were analyzed by Western blotting, probed with antibodies against GFP, Talin, Abi1, Nckap1, Cyfip1, Wasf1 and 14-3-3. (***E***) Flow cytometry analysis of GFP and integrin β1 expression in GE11 ^tetON^ ^β1^ cells stably expressing the indicated construct.

